# Structural Integrity of RyR2 Clusters Controls Cardiac Calcium Leak

**DOI:** 10.1101/2025.11.07.687312

**Authors:** Andrew Noren, Yohannes Shiferaw

## Abstract

Calcium (Ca) leak from the sarcoplasmic reticulum is a key contributor to cardiac arrhythmias, yet the structural mechanisms that regulate spontaneous Ca release from ryanodine receptor type 2 (RyR2) clusters remain poorly understood. To investigate how cluster architecture controls Ca leak, we developed a computational model in which each RyR2 channel is composed of four interacting subunits and embedded within spatially organized clusters. This framework captures both cooperative gating within individual channels and coupling between neighboring channels. Our simulations reveal that the timing of spontaneous Ca sparks is exponentially dependent on the structural integrity of the RyR2 cluster. This exponential sensitivity means that even small disruptions in cluster structural integrity, such as partial fragmentation, can lead to a 100-1000 times increase in spontaneous Ca spark frequency. These findings identify cluster structural integrity as a powerful control mechanism for Ca leak, and represent a promising therapeutic target for restoring Ca homeostasis in cardiac myocytes.

## Introduction

Calcium (Ca) signaling is fundamental to cardiac muscle contraction, with tightly regulated intracellular Ca release essential for effective excitation-contraction coupling(1). The RyR2, a large Ca release channel located on the sarcoplasmic reticulum (SR) membrane, plays a central role in this process. RyR2 is activated through calcium-induced calcium release (CICR), in which a small influx of Ca via L-type Ca channels (LCCs) during the cardiac action potential triggers a much larger release of Ca from the SR. Recent advances in super-resolution imaging techniques, have provided unprecedented insight into the spatial organization of RyR2 within cardiomyocytes (2–8) These methods have revealed that RyR2 channels form distinct, heterogeneous clusters whose size, density, and arrangement are critical determinants of Ca signaling dynamics. This detailed structural information now provides a framework for investigating how the organization of RyR2 clusters contributes to normal cardiac physiology, and how its disruption may lead to arrhythmogenesis(9–14).

New imaging techniques have revealed that RyR2 clusters are not static structures, but instead show dynamic organization that can change significantly in disease (6,10,15). In healthy cardiomyocytes, RyR2 channels are arranged in compact organized clusters at dyads. In disease, however, this organization becomes disrupted. A common observation is cluster fragmentation, where large RyR2 assemblies break into smaller, irregularly shaped groupings. This reduces the number of channels per cluster and alters their spatial arrangement, with important consequences for Ca signaling. In an elegant study Sheard et al. (2) used enhanced expansion microscopy to visualize this remodeling in three dimensions, revealing that fragmented RyR2 clusters often exhibit a frayed appearance. In these clusters, RyR2 channels remain partially anchored near the center of the cluster, where a structural protein called junctophilin-2 (JPH2) helps tether them to the membrane, but many RyRs extend outward in a disorganized, loosely connected pattern. This “fraying” suggests a loss of structural integrity in the regions where JPH2 is absent, marking the early breakdown of the tightly organized release sites essential for normal Ca signaling. In heart failure, similar changes are observed. Super-resolution imaging studies, such as those by Kolstad et al. (6), have shown that RyR2 clusters become dispersed, forming smaller and more loosely organized sub-clusters. Also, in persistent atrial fibrillation, Macquaide et al. (11) demonstrated that RyR2 cluster fragmentation results in more numerous, closely spaced clusters within Ca release units, accompanied by a greater than 50% increase in spontaneous Ca spark frequency. These structural changes are closely linked to key functional defects in heart failure, including elevated diastolic Ca leak, reduced contractile strength, and a greater risk of arrhythmias(13,16,17).

Computational models have provided important insights into how RyR2 spatial organization affects calcium signaling. Recent studies have demonstrated that RyR2 interactions via Ca diffusion are critical for coordinating calcium release (18), with network connectivity affecting whole-cell calcium oscillations (19) and cluster geometry influencing calcium spark properties(20,21). These findings establish that spatial effects operate through multiple mechanisms, particularly diffusive calcium coupling between channels. However, existing models have not incorporated direct subunit coupling between adjacent RyR2 channels within clusters. Such coupling arises from the physical contacts between neighboring channels revealed by cryo-EM studies (22) and represents a distinct mechanism through which cluster connectivity may regulate calcium release. In this study, we model a cluster of RyR2 channels as an array of interacting tetramers. Each RyR2 channel is composed of four subunits, and channels within the cluster are arranged with specific geometrical contacts that reflect their structural organization. We analyze the stochastic dynamics of this system to determine how the arrangement of neighboring channels influences the behavior of the cluster. Our focus is to understand how cluster architecture controls the timing of spontaneous Ca sparks. A central finding is that, at resting Ca concentrations, the frequency of spontaneous Ca sparks is exponentially sensitive to the structural integrity of the cluster. In this regime, small changes in arrangement, such as increased spacing, reduced connectivity, or fragmentation into sub-clusters, can increase the frequency of spontaneous Ca sparks by several orders of magnitude. This result highlights the crucial role of intact cluster architecture in maintaining coordinated channel closure at rest. We further show that frayed clusters are particularly vulnerable, since small peripheral groups of RyR2s can recruit larger assemblies through diffusive Ca coupling. Together, these results identify cluster architecture as a dominant control mechanism for Ca leak under diastolic conditions, linking cluster integrity to pathological Ca handling.

## Methods

### Computational Model of RyR2 tetramer

The RyR2 channel is a tetramer composed of four identical subunits that gate a shared central pore. Structural studies show that these subunits are tightly packed and physically interact at their interfaces(22,23). Motivated by these observations and by our earlier modeling work(24), we describe a simplified RyR2 model in which each subunit can exist in one of two conformational states. A central assumption of the model is that adjacent subunits interact energetically, such that conformational mismatches are penalized. In particular, if a subunit transitions from closed to open while its neighbors remain closed, the resulting mismatch produces an energetic cost. This is implemented by modifying the transition rate so that the opening rate of a closed subunit is reduced by a multiplicative factor when neighbors are closed and increased when neighbors are open. An additional assumption is that a channel is considered functionally open when three or more subunits are in the open state. This requirement was established in our previous study (24), where we demonstrated that RyR2 channels must exhibit cooperative gating to remain reliably shut at diastolic Ca concentrations. Without this constraint, channels cannot maintain stable closure during rest.

To apply this model, we represent the RyR2 channel as four subunits labeled *s*_*i*_, with *i* = 1.2.3.4. and periodic boundary conditions. Each subunit can be in one of two states: closed (*s*_*i*_ = −1) or open (*s*_*i*_ = +1). This configuration ensures that every subunit interacts with its two immediate neighbors, consistent with the geometry of the RyR2 channel. Transitions between subunit states are governed by rates that depend on the conformational states of neighboring subunits. The forward (opening) and backward (closing) transition rates for subunit *i* are given by

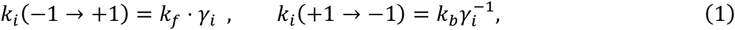

where *k*_*f*_ and *k*_*b*_ are the intrinsic forward and backward rates in the absence of coupling. The factor *γ*_*i*_ captures the influence of neighboring subunits and is defined as:

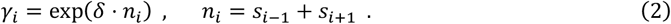

Here, *δ* = *βJ* is a dimensionless parameter representing the strength of coupling between channels, and where *J* is the energy penalty. Also, *β* = 1/*k*_*B*_*T* where *k*_*B*_ is Boltzman’s constant, and *T* is the temperature. These local interaction rules form the basis of a thermodynamically consistent model that accounts for cooperative interactions within individual RyR2 channels.

### Computational model of an RyR2 cluster

Recent Cryo-EM imaging studies due to Cabra et al. (22) reveals that RyR2 channels physically interact at their interfaces through specific molecular contacts between adjacent channels. Their high-resolution structural analysis identifies two distinct types of inter-channel arrangements: an “adjoining” configuration (Figure 1A) and an “oblique” configuration (Figure 1B). In the adjoining arrangement, channels are aligned in regular rows and columns, with horizontal neighbors coupling through their edge-to-edge contacts. In the oblique arrangement (Figure 1B), channels adopt a staggered geometry with interactions between subunits 1-3 and 2-4 only. Cabra et al. observed that native RyR2 clusters typically form disordered structures that represent combinations of these two geometric arrangements rather than uniform arrays of a single type. To model these experimentally observed arrangements, we will consider cluster models based on both interaction types.

**Figure 1.**
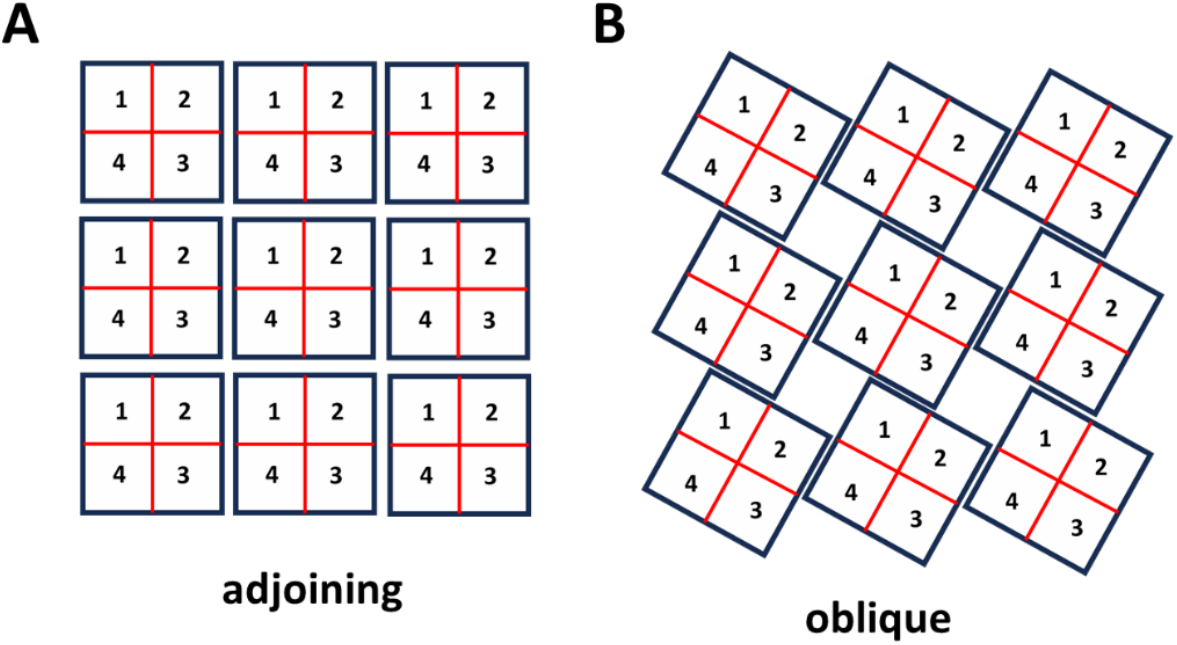
Architecture of RyR2 clusters. **(A)** *Adjoining configuration*. RyR2 tetramers are positioned in a side-by-side array, with neighboring channels aligned in rows and columns. In this arrangement, adjacent channels make contact along their lateral surfaces, consistent with the “side-by-side” geometry observed in cardiac cells. **(B)** *Oblique configuration*. RyR2 tetramers are connected in an alternating “checkerboard” pattern. This arrangement corresponds to the oblique interaction identified in structural studies, where subunits from diagonally offset channels (1–3 and 2–4) form the interface.

To model heterogeneous arrays, we denote 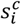 as the state of subunit *i* in channel *c*. Similarly we will denote 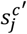 and 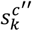 to be the state of subunits *j* and *k* in neighboring channels *c*′ and *c*″ respectively. The coupling factor for subunit *i* in channel *c* is given by

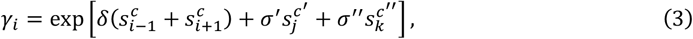

where *σ*′ and *σ*″ represents the subunit interaction strength between subunit *i* in channel *c* and subunits *j* and *k* in neighboring channels *c*′ and *c*″. For simplicity, we will assume *σ*′ = *σ*″ = *σ* in this study. However, it is likely that in realistic scenarios these parameters may differ due to variations in channel properties. With this formulation the transition rates have the same form as in Eq. (1) and remain consistent with the principle of detailed balance.

To model inter-RyR2 interactions within a cluster, we arrange RyR2 channels either with the oblique or adjoining configurations shown in Figure 1. Each channel occupies a grid position labeled by coordinates (*n*_*x*_.*n*_*y*_), and consists of four subunits labeled *s*_*i*_. Each subunit is uniquely identified by the triplet (*n*_*x*_.*n*_*y*_.*s*_*i*_), which denotes the lattice position and the state of the tetramer. The model incorporates two distinct types of coupling interactions. First, intra-channel coupling occurs between adjacent subunits within the same RyR2 tetramer, as described in Eq. (2). Second, inter-channel coupling links specific subunits from neighbouring channels according to the geometric constraints of each configuration. In the adjoining arrangement, subunits form connections with two nearest-neighbour channels, reflecting the side-by-side contact geometry. In the oblique arrangement, each subunit couples to only one nearest neighbour, corresponding to the staggered interaction pattern. To capture more realistic cluster arrangements, we will also consider disordered configurations that reflect the structural heterogeneity observed in cardiac myocytes. This computational framework will allow us to systematically investigate how the loss of structural integrity affects Ca release dynamics.

### Model of Ca regulation of RyR2 cluster dynamics

Super-resolution imaging studies reveal that RyR2 cluster sizes span a wide range. The most recent work by Hou et al. (25) reports many small clusters containing roughly 20–50 channels (spanning approximately 150–250 nm in diameter), while earlier studies by Galice and colleagues (26) observed larger aggregates that could extend to several hundred RyR2s, with typical averages closer to 50–100 channels (spanning approximately 250–350 nm in diameter). Although the absolute estimates vary across techniques, the consistent finding is that ventricular myocytes contain a heterogeneous population of clusters, ranging from solitary channels and small groups to large assemblies. The functional behaviour of these clusters is strongly shaped by Ca diffusion in the narrow dyadic cleft. Ca diffuses rapidly in the cytosol, with an effective diffusion coefficient in the range *D* ≈ 100 − 500(*μm*)^2^/*s* (27,28). This wide range is due to the presence of multiple buffers and the structural complexity of the intracellular space. Given an RyR2 open time of roughly *τ* ~ 1*ms*, the expected diffusion length (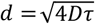) is in the range 600 − 1000*nm*. This length scale exceeds the size of even the largest RyR2 clusters (≤ 400*nm*), implying that Ca released from a single open channel rapidly equilibrates across the entire cluster. As a result, RyR2s within a cluster are functionally coupled through the shared local Ca signal, independently of direct molecular contact.

Since diffusion is much faster than channel gating, we invoke the rapid diffusion approximation, which assumes Ca is uniform within the dyad. Under this approximation, the local Ca concentration is given by

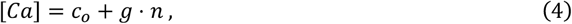

where *c*_0_ is the diastolic Ca concentration in the bulk cytosol, *n* represents the number of open RyR2 channels within the cluster, and *g* represents the Ca concentration increase in the dyadic cleft due to a single open RyR2 channel(27,29,30). Several computational studies of CICR have argued that [Ca] must reach relatively high concentrations in the dyadic junction. Cannell et al. (18) argued that minimum dyadic [Ca] of ~50*μM* was required to maintain CICR in rat ventricular myocytes, while Shen et al. (31) used half-maximal RyR2 activation concentrations (*K*_*d*_) ranging from 25 − 45*μM* in their models of RyR cluster dispersion during β-adrenergic stimulation. In this study we will set *g* = 25*μM*, so that 1-3 open channels produce dyadic [Ca] in the range of 25-75 μM, consistent with the concentrations required to support CICR in these studies.

To determine the transition rates in our model, we assume first-order kinetics with respect to cytosolic Ca. Specifically, the forward rate for subunit opening is given by 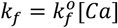, where 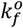 is a binding rate constant and [*Ca*] is the local Ca concentration. We set 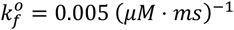 to ensure that RyR2 channels remain predominantly closed at a diastolic concentratonof [*Ca*] ~ 0.1*μM*. This choice is consistent with experimental observations, see for example Shen et al. (31), showing that spontaneous Ca sparks are extremely rare at baseline (~ 0.1 *sparks*⁄100*μm*/*s*). Although this forward rate is substantially lower than binding rates measured in bilayer recordings (32,33), the difference reflects the highly regulated environment of intact cells. In particular, physiological *Mg*^2+^ occupies the Ca-activation sites at resting [Ca] and must be displaced by Ca to allow channel opening, effectively reducing the Ca sensitivity and stabilizing the closed state compared to isolated channels in bilayers(34–36). The backward rate for subunit closing is Ca-independent and fixed at *k*_*b*_ = (*ms*)^−1^ (33), which corresponds to an average open time of approximately 3*ms*. Subunit cooperativity is captured by a dimensionless mismatch penalty *δ* = 0.5, which stabilizes the closed conformation at rest without imposing such a large energetic cost that the channel becomes unresponsive at elevated Ca. A pore conducts only when at least three of the four subunits are open, a majority rule that preserves observed closed-time statistics and prevents spurious openings(24). Collectively, these parameters produce physiologically plausible single-channel kinetics and cluster-level behavior across the physiological Ca range.

### Stochastic simulation algorithm

The Gillespie algorithm is implemented as a stochastic simulation method to model the exact temporal evolution of RyR2 channel gating within clusters(37). At each time step, the algorithm calculates the total reaction rate across all subunits in the system based on their current states and local coupling environments. The time to the next reaction is drawn from an exponential distribution with parameter equal to the total rate, ensuring proper stochastic timing. A specific reaction is then selected using weighted random selection proportional to individual subunit rates, after which the chosen subunit transitions between closed and open states. This process continues iteratively, with the system state analyzed after each reaction to count the number of open channels.

## Results

### Measuring the timing of a spontaneous Ca spark

To study the stochastic dynamics of RyR2 clusters, we first simulated the time evolution of a 10 × 10 array of RyR2 channels using our stochastic simulation algorithm. To monitor the cluster, we keep track of the number of open RyR2 channels, denoted as *n*_*o*_, which is defined as the number of channels where 3 or more subunits are in the open state. Figure 2 shows *n*_*o*_ as function of time for three simulation runs where the cytosolic Ca concentration is fixed at *c*_*o*_ = 6.0*μM*. Each independent simulation run is shown as a red, black, and blue line. In these runs the cluster stays mostly closed and occasionally one or two channels transition to the open state. After a waiting period a large fluctuation pushes *n*_*o*_ above a critical threshold, leading to a rapid cooperative avtivation of the entire cluster. Once this transition occurs, the number of open channels grows explosively from a few (*n*_*o*_ ~ 0 − 1) to nearly all the channels in the cluster (*n*_*o*_ ~ 100). This behaviour arises because the local Ca concentration is dependent on *n*_*o*_ according to Eq. (4). Thus, the opening rate of channels increases in a strongly nonlinear fashion due to subunit cooperativity within and between RyR2 channels.

**Figure 2.**
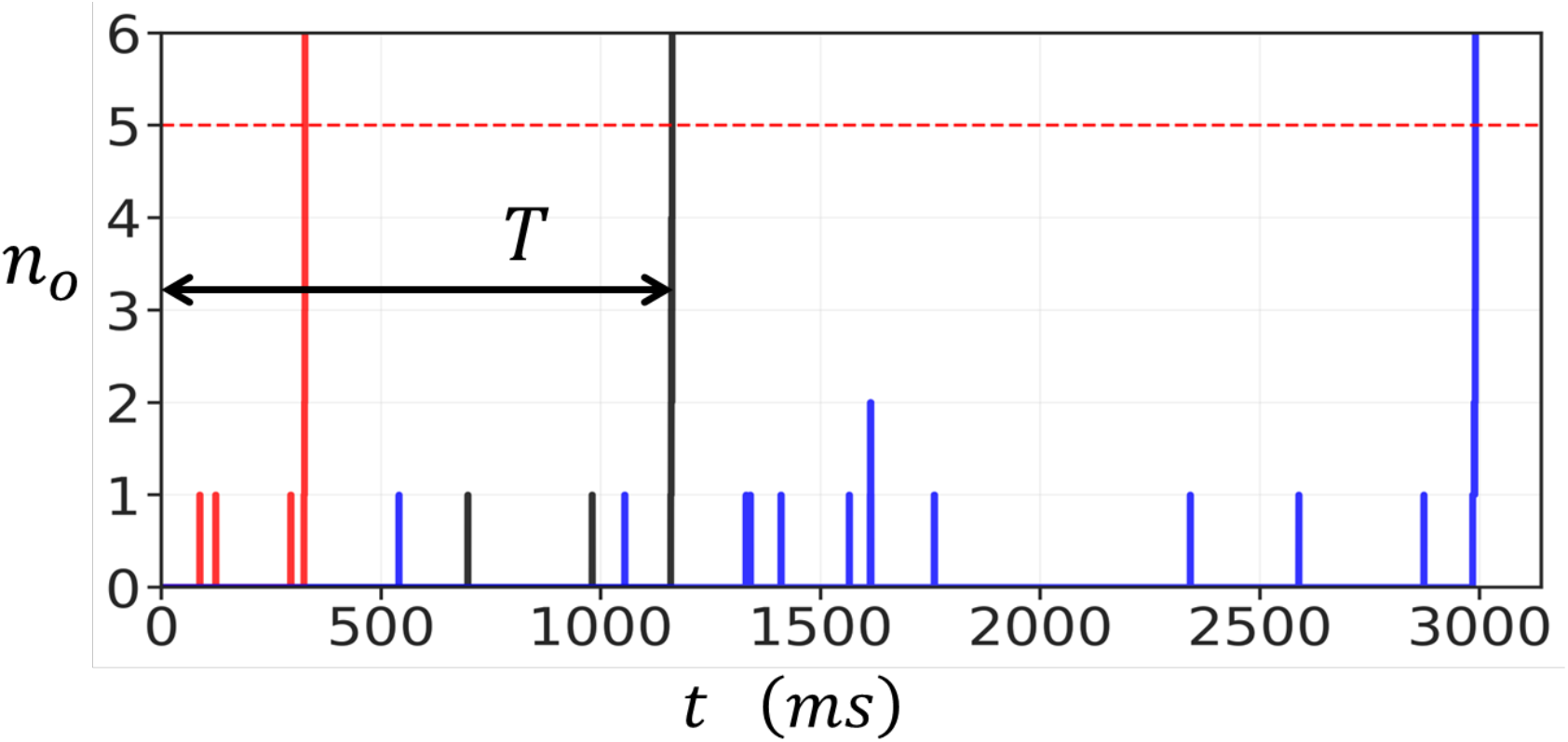
Stochastic activation of a 10 × 10 RyR2 cluster connected in the oblique configuration. The cytosolic Ca concentration is fixed at *c*_*o*_ = 6.0*μM*. Three independent simulation runs are shown corresponding to the red, black, and blue lines. The activation time *T* (for the simulation shown in black) is the time when the channels are all closed, to when *n*_*o*_ exceeds *n*_*c*_ = 5. The parameters used in this simulation are 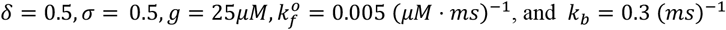.

The rapid increase in *n*_*o*_ corresponds to the initiation of a Ca spark. Since the excitation occurred because of stochastic fluctuations, we will refer to these events as spontaneous Ca sparks. After activation, RyR2 channels proceed to inactivate and return to the closed state. However, this inactivation process occurs on a much shorter timescale than the waiting time to a spontaneous Ca spark and is not modelled in this study. To quantify the spontaneous Ca spark initiation process, we define the activation time *T*, shown in Figure 2 for the simulation run corresponding to the black line, as the time required for the cluster to undergo the initial transition from the closed state to a fully activated cluster. Specifically, *T* is measured from *t* = 0, when all RyR2 channels in the cluster are initially closed (all subunits are in the state *s*_*i*_ = −1), to the moment when *n*_*o*_ exceeds a critical threshold *n*_*c*_. In general, *n*_*c*_ should be picked to be larger than a typical fluctuation, and in this study we will use *n*_*c*_ = 5. However, we note here that our results are independent of *n*_*c*_ providing *n*_*c*_ is larger than the baseline fluctuations. For example, in the simulation shown in Figure 2, any *n*_*c*_ ≥ 5 gives the same waiting time since once *n*_*c*_ = 5 is reached then excitation proceeds with high likelihood. This approach provides a consistent measure of the waiting time before spontaneous Ca spark initiation, which determines the average frequency of spontaneous Ca sparks in the cell.

### Control of Spontaneous Spark Timing by Diastolic Ca and Cluster Coupling

We next investigated how the mean waiting time *T* to a spontaneous Ca spark depends on the dyadic Ca concentration *c*_*o*_. This concentration is typically *c*_*o*_ ~ 0.1*μM* during rest, but will increase substantially when there is a local opening of a nearby LCC channel. Figure 3A shows the relationship between *T* and *c*_*o*_ for the oblique configuration of the cluster. Here, we vary *c*_*o*_ from 5.0*μM* to 120*μM*, which represents the typical Ca concentrations at a dyadic junction that occurs during LCC activation. Three values of the inter-subunit coupling parameter *σ* are examined: *σ* = 0 (no coupling), *σ* = 0.5 (moderate coupling), and *σ* = 1 (strong coupling). The results show that *T* changes exponentially with the concentration *c*_*o*_. At high diastolic Ca (*c*_*o*_ ~ 100*μM*) all three cases have comparable firing times in the range 0.5*ms* − 2*ms*. In contrast, at lower Ca concentrations the waiting time increases exponentially with decreasing *c*_*o*_. In particular, when subunit interactions between channels is engaged, with *σ* = 1.0, the waiting time increases from ~ 2*ms* at *c*_*o*_ ~ 120*μM*, to ~ 10^5^*ms* at *c*_*o*_ ~ 5.0*μM*. Figure 3B shows analogous results for the adjoining configuration. The qualitative features are the same as for the oblique case, but the dependence on *σ* is more pronounced, since each subunit has two contacts rather than just one. In Figure 3C and 3D we compute *T*, for both the oblique and adjoining configurations respectively, for cluster sizes of 5 × 5, 10 × 10, and 20 × 20. These results shows that the dependence of *T* on *c*_*o*_ is similar across the wide range of cluster sizes that are expected to be found in cardiac myocytes.

**Figure 3.**
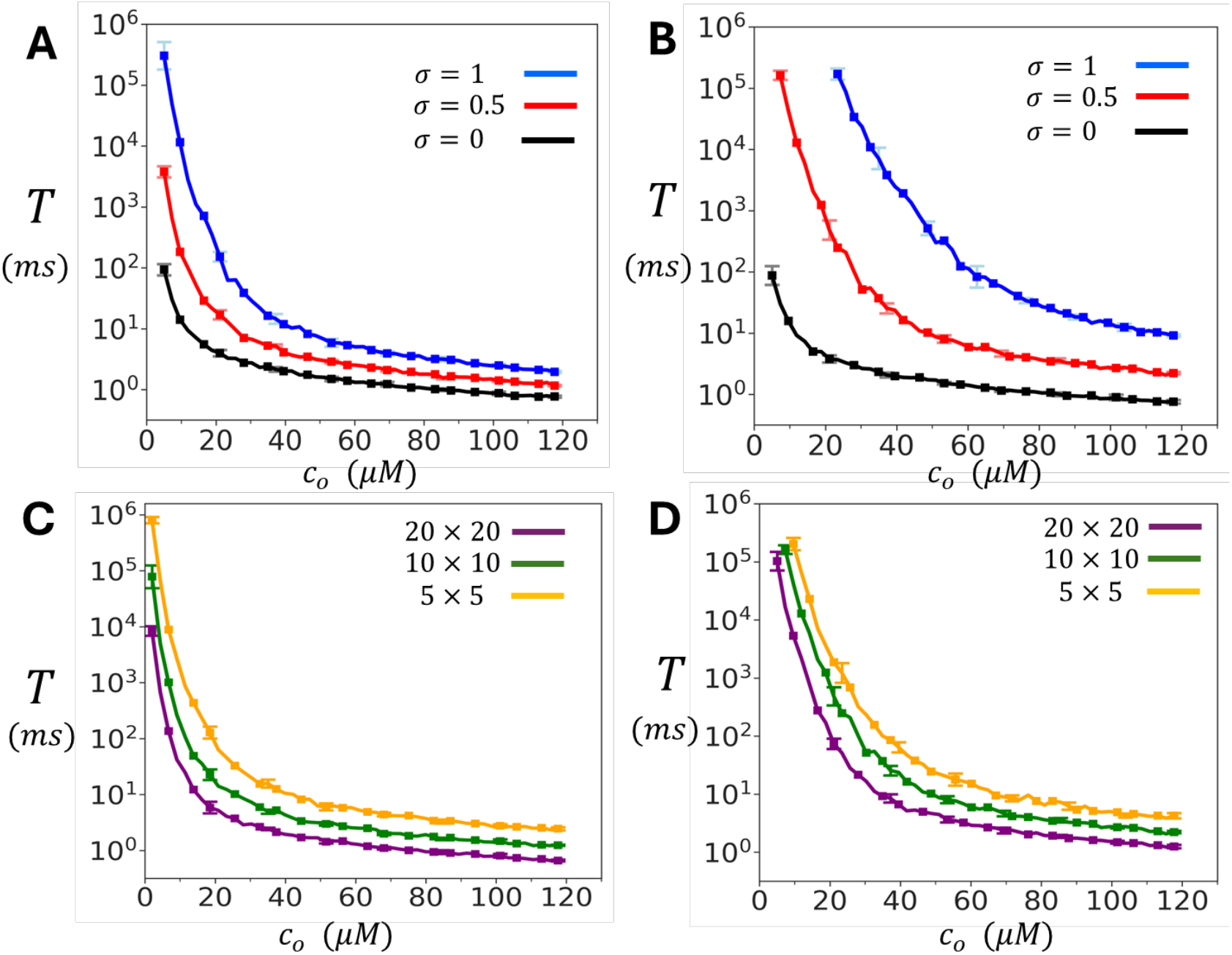
Dependence of spontaneous spark timing on diastolic Ca *c*_*o*_. **(A)** Mean waiting time *T* to a spontaneous Ca spark as a function of diastolic Ca concentration *c*_*o*_ for the oblique cluster configuration. *T* is computed by averaging over 50 independent simulation runs. Error bars shown are computed as the standard deviation of 5 independent sets of 10 simulation runs each. **(B)** Same simulation as in **A** but with RyR2 channels arranged in the adjoining configuration. **(C)** Same simulations as in **A** but with cluster sizes of 5 × 5. 10 × 10. and 20 × 20. **(D)** Same simulations as in **B** with the indicated cluster sizes. For all simulations in C and D we have fixed *σ* = 0.5.

To further explore the role of coupling, we examined the dependence of *T* on *σ* at two fixed Ca concentrations. Figure 4A shows results at a resting concentration of *c*_*o*_ = 5.0*μM*. In this regime, the mean waiting time measured for both the oblique and adjoining configurations grows almost three orders of magnitude as *σ* is increased. In contrast, at a diastolic Ca concentration of *c*_*o*_ = 100*μM*, shown in Figure 4B, *T* increases only modestly over a few milliseconds. Together, these findings identify inter-channel coupling as a powerful mechanism for tuning cluster stability at low Ca concentrations. At high Ca, clusters activate rapidly and reliably, with little dependence on coupling strength. At low Ca, however, even modest increases in *σ* dramatically stabilize the closed state by exponentially prolonging the waiting time *T*. This dual behaviour ensures that RyR2 clusters remain quiescent at rest, while still capable of fast activation when triggered during excitation– contraction coupling.

**Figure 4.**
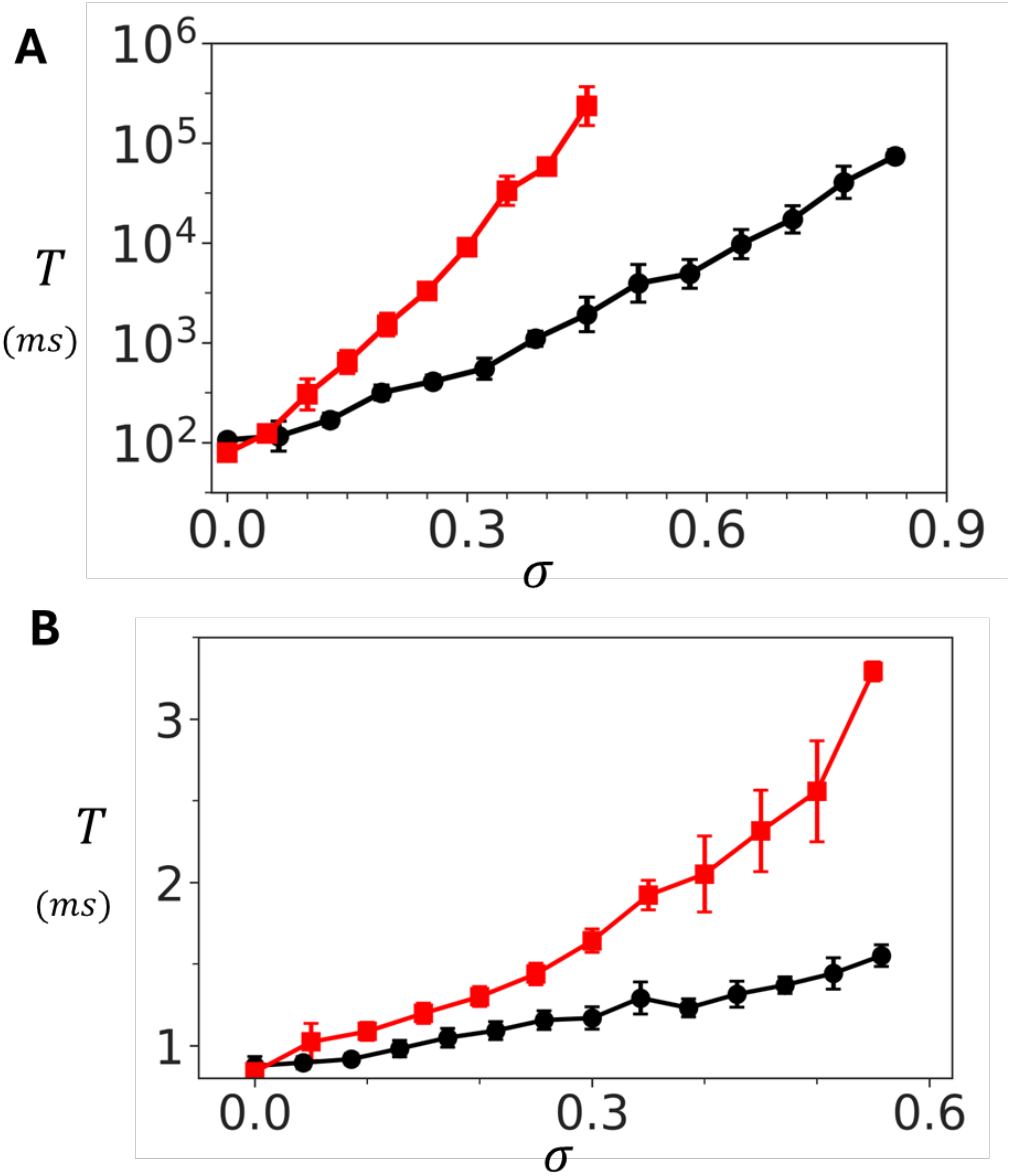
Dependence of spontaneous Ca spark timing on inter-subunit coupling strength. **(A)** Dependence of *T* on *σ* at *c*_*o*_ = 5.0*μM* for a 10 × 10 cluster. Red (black) line corresponds to the adjoining (oblique) configuration. **(B)** Dependence of *T* on *σ* at high Ca (*c*_*o*_ = 100*μM*). The waiting time increases only modestly, indicating weak coupling effects at elevated Ca. Points plotted are averaged over 50 samples and the error bar error bars shown are estimated by computing the standard deviation of 5 independent sets of 10 runs each.

### Modeling heterogeneous clusters

Experimental studies have revealed that RyR2 clusters undergo fractionation and lose their structural integrity in diseased states. To investigate the functional consequences of cluster fractionation, we developed a computational model that simulates the progressive disruption of RyR2 cluster connectivity. Our approach implements stochastic bond breaking between adjacent RyR2 channels within the cluster. Specifically, we assign a probability *p* that a connection between two nearest-neighbor subunits remains intact. The parameter *p* serves as a measure of cluster integrity, where *p* = 1 corresponds to a fully connected cluster with all inter-channel connections preserved, while *p* = 0 represents a completely dissociated cluster with all connections broken. For each value of *p*, we generate an ensemble of connectivity matrices that define the network topology and compute the average *T* over the ensemble. In Figure 5, we show the relationship between cluster connectivity and spontaneous activation by plotting *T* versus the connection probability *p* at a Ca concentration of *c*_*o*_ = 2.0*μM*. The results show that the waiting time *T* grows exponentially with increasing connection probability *p*. In particular, for the case of the adjoining configuration, we find that an uncoupled cluster of 10 × 10 RyR2s has a mean firing time of *T* ~ 10^3^*ms*. However, as *p* is increased to just *p* ~ 0.5, the waiting time rises to *T* ~ 10^6^*ms*. This, result shows that the spontaneous Ca spark frequency is exponentially sensitive to the probability that neighboring RyR2s are connected.

**Figure 5.**
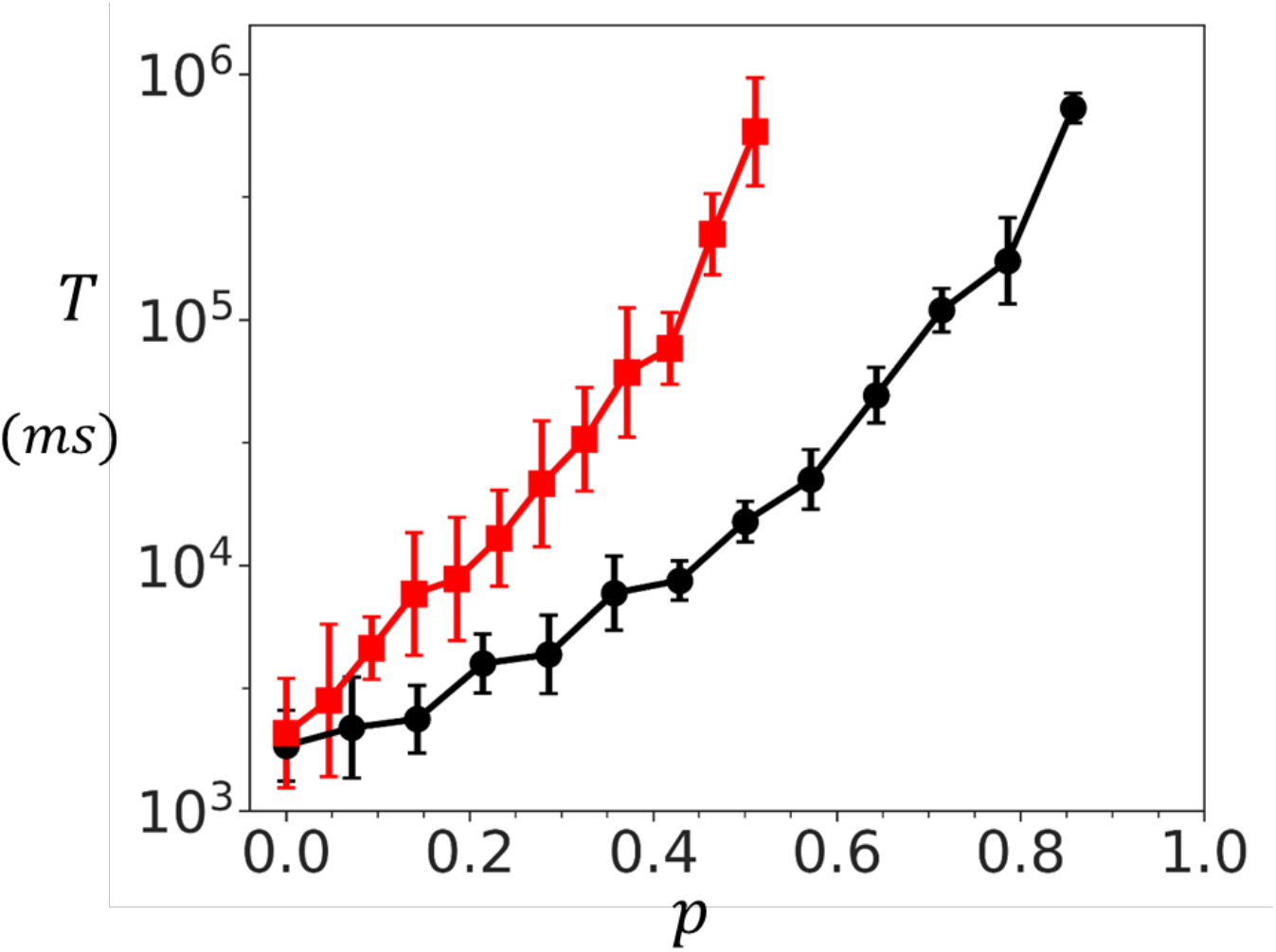
Dependence of spontaneous spark timing on cluster connectivity. Mean waiting time *T* to a spontaneous Ca spark as function of the connection probability *p*. In this simulation the connectivity strength is set to *σ* = 1.0 and *c*_*o*_ = 2.0*μM*. Red line (black line) denotes the adjoining (oblique) cluster configurations of size 10 × 10. Here, *T* is computed by averaging over 50 independent simulations, and the error bars are computed as the standard deviation 5 sets of 10 simulation runs each.

Experimental studies reveal that RyR2 clusters within the junctional sarcoplasmic reticulum exhibit highly heterogeneous spatial arrangements. To investigate how this structural diversity affects functional properties, we have also developed a preferential attachment growth model that generates realistic cluster morphologies. The preferential attachment algorithm sequentially places RyR2 channels on a two-dimensional lattice. For each new channel placement, the probability of selecting an empty site (*i.j*) is proportional to (1 + *n*_*e*_)^α^. where *n*_*e*_ represents the number of occupied nearest-neighbor sites and the exponent *α* is referred to as the clustering parameter. This formulation ensures that sites adjacent to existing channels are preferentially selected, with the strength of this preference controlled by *α*. When *α* = 0, channel placement is uniform and random, while increasing values promote increasingly compact cluster formation.

Figure 6A demonstrates the range of cluster structures generated by varying the clustering parameter *α*. At *α* = 0, RyR2 channels are scattered randomly on the grid. At intermediate values (*α* = 5) more cohesive structures start to from, while retaining some fragmentation. A high clustering parameter (*α* = 8) generates compact arrangements with a few dispersed channels. To examine how cluster structural heterogeneity affects spontaneous Ca spark dynamics, we applied the preferential attachment algorithm by placing 50 RyR2 channels on a 20×20 lattice domain. Given that individual RyR2 channels are approximately 30*nm* × 30*nm* in size (22), this domain corresponds to approximately 600*nm* in diameter, roughly the spatial extent over which Ca is strongly diffusively coupled. For each value of the clustering parameter *α*, we generated 50 independent cluster configurations. Once the configuration is generated the channels are coupled according to the oblique architecture. We then average the mean first passage time *T* over all configurations. Figure 6B shows *T* vs *α* at a fixed *c*_*o*_ = 2.5*μM*. The results show that *T* increases substantially as *α* is varied in the range 0 to 10. At low values (*α* ≈ 0), corresponding to dispersed clusters with weak inter-channel connectivity, the mean waiting time is approximately 5 × 10^3^*ms*. As *α* increases, producing tightly organized clusters, *T* increases to approximately 10^5^*ms* for the oblique configuration, and 10^6^ for the adjoining configuration. This effect saturates above *α* ≈ 7 since all clusters generated for larger *α* are close to fully compact and yield the same *T* as the fully connected square lattice. This exponential relationship, in the range 0 to 7, demonstrates that cluster compactness is a critical determinant of spontaneous Ca release dynamics, with tightly organized structures providing exponentially greater stability during diastole.

**Figure 6.**
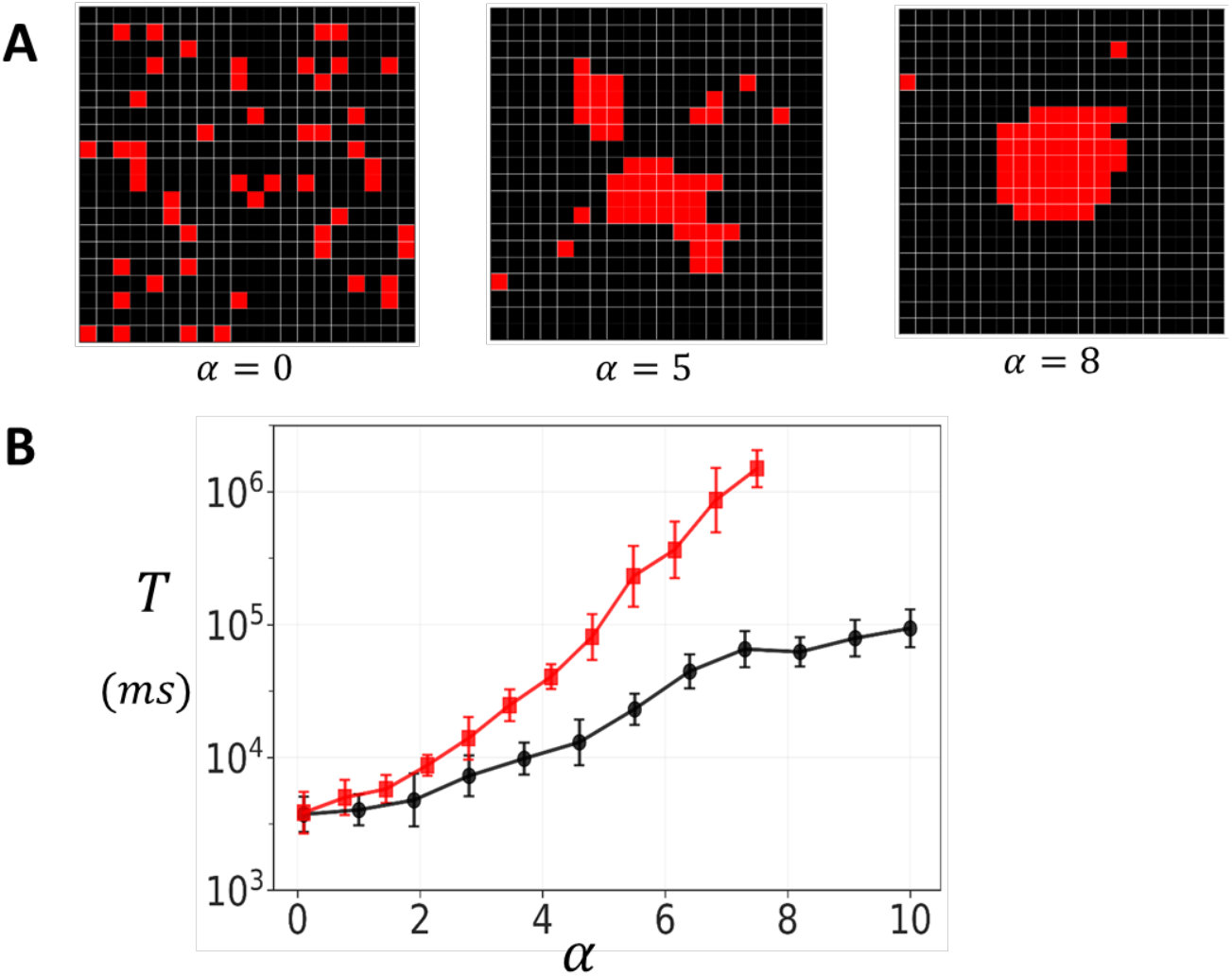
Spark timing in heterogeneous clusters. **(A)** Representative cluster morphologies generated by varying the clustering parameter *α* in the preferential attachment algorithm. Low values yield dispersed clusters with irregular boundaries, intermediate values produce partially cohesive structures, and high values generate compact, dense aggregates. In this simulation we place 50 RyR2 channels on a grid of size 20 × 20. Neighboring channels are connected according to the oblique geometry. **(B)** Dependence of the mean waiting time *T* for spontaneous Ca spark initiation on the clustering parameter. Red and black line corresponds to *σ* = 1 and *σ* = 0.5 respectively. The waiting time *T* is computed by averaging over 50 independent simulation runs. For each simulation run a different cluster arrangement is generated using the preferential growth algorithm. Here, we set *c*_*o*_ = 2.5*μM*, and error bars are computed using 5 sets of 10 simulation runs.

## Discussion

### Inter-subunit coupling controls cluster response to Ca

In this study, we developed a computational model of an RyR2 cluster in which each channel is composed of four interacting subunits, with channels arranged in a configuration consistent with structures observed in cryo-EM and super-resolution imaging. Our goal was to understand how inter-subunit cooperativity influences Ca leak in cardiac cells. The first result of the model is that the cluster exhibits a sharp contrast in behavior across different Ca concentration regimes. At Ca concentrations larger than 100*μM* RyR2 clusters respond rapidly to Ca signals on sub-millisecond timescales. This fast response is largely insensitive to the strength of channel-to-channel coupling. However, at lower Ca concentrations (below ~ 5*μM*), the system behaves very differently. In this case, the mean waiting time to a spontaneous Ca spark becomes exponentially sensitive to the strength of subunit coupling. Specifically, at a Ca concentration of 5.0*μM*, increasing the coupling strength *σ* between RyR2 subunits from 0 to 0.5 (for the adjoining arrangement), leads to an almost 4 orders of magnitude increase in the mean waiting time. This behavior highlights the functional significance of inter-subunit coupling: it preserves fast responsiveness to LCC openings while ensuring cluster stability at lower Ca concentrations. An important consequence of this finding is that diastolic Ca leak, which is determined by how often spontaneous Ca sparks occur, will be exponentially sensitive to the coupling between RyR2 within clusters. Therefore, even small changes in coupling strength can produce orders-of-magnitude differences in leak rate. These results highlights the critical role of RyR2 coupling in regulating Ca cycling homeostasis.

### Ca leak is exponentially sensitive to RyR2 cluster integrity

Experimental evidence from cryo-EM and super-resolution microscopy shows that RyR2 clusters in cardiac cells are highly irregular, forming heterogeneous structures rather than perfect lattices. To examine how this heterogeneity shapes Ca signaling, we extended our modeling framework to include disordered cluster geometries and systematically varied their structural integrity. In one approach, we introduced a coupling parameter *p*, defined as the probability that a subunit remains linked to its nearest neighbor. This parameter represents the progressive loss of inter-subunit connectivity. As *p* decreased from 1 (fully intact) to 0 (fully fragmented), the mean waiting time to a spontaneous Ca spark shortened by approximately 3 orders of magnitude. These results shows that Ca spark frequency is exponentially sensitive to even small disruptions in cluster connectivity. In parallel, we used a preferential attachment algorithm to generate clusters with heterogeneous shapes resembling those observed experimentally. Here, the clustering parameter *α* controlled the likelihood that new channels attached to existing groups, thereby tuning the degree of structural cohesion. Increasing *α* produced a substantial increase in spark waiting time, demonstrating that more tightly connected clusters are exponentially more stable at lower Ca concentrations. Together, these results reveal that heterogeneity in RyR2 cluster organization plays a central role in regulating the timing of spontaneous Ca sparks.

### Ca leak in heart failure is dependent on RyR2 structural integrity

Recent high-resolution imaging studies have revealed that RyR2 clusters lose their structural integrity in heart failure, with important functional consequences for Ca handling. Sheard et al. (2) used enhanced expansion microscopy to demonstrate that RyR2 clusters in failing myocytes frequently exhibit a frayed appearance, where small groups of channels detach from the main cluster, particularly in regions depleted of the structural protein junctophilin-2. Similarly, Kolstad et al. (6) showed that post-infarction heart failure is characterized by RyR2 cluster dispersion and fragmentation, resulting in smaller, more numerous cluster fragments with reduced inter-channel connectivity. Our computational modelling provides a mechanistic explanation for why these structural changes change the Ca leak rate. The key insight from our model is that spontaneous Ca spark frequency exhibits exponential sensitivity to cluster structural integrity. Mechanistically, intact inter-subunit coupling within the cluster provides stabilizing interactions that maintain channels in the closed state during diastole through cooperative inhibition. When coupling is weakened by fragmentation, this stabilizing effect is lost, making the cluster far more susceptible to stochastic activation. The key new insight in this study is that this relationship is exponential, and explains why even modest cluster remodeling, such as partial fragmentation or fraying, can produce the orders of magnitude increases in diastolic Ca leak observed in heart failure. Thus, cluster architectural integrity emerges as a critical control mechanism that drives Ca leak.

### The Role of Diffusion in RyR2 Cluster Dynamics

Ca diffusion plays a crucial role in determining the functional coupling between RyR2 channels within the dyadic cleft. With a diffusion coefficient of approximately 100 − 500 (*μm*)^2^/*s* and an RyR2 open time of roughly 1*ms*, Ca diffuses roughly 600*nm* before channel closure. Since this diffusion length exceeds the diameter of even the largest RyR2 clusters (≤ 400*nm*), Ca released from any open channel rapidly equilibrates across the entire dyadic space, creating strong functional coupling through the shared Ca microdomain. Importantly, when clusters become frayed or fragmented into separate pieces, the disjoint subclusters often remain within the same Ca microdomain and are therefore still functionally coupled through diffusion. However, these separated channels lose the stabilizing nearest-neighbor interactions that maintain closure during diastole. A small detached subcluster can thus spontaneously open without the energetic penalty of intact inter-channel coupling yet still elevate local Ca sufficiently to recruit the larger cluster. This asymmetry between lost stabilization and preserved diffusive coupling explains why frayed clusters exhibit exponentially faster spontaneous Ca spark frequencies.

### Cluster integrity as a therapeutic target

The exponential sensitivity of Ca leak to RyR2 coupling revealed in our study carries significant therapeutic implications. Because spark frequency rises exponentially as coupling weakens, even subtle structural disruptions can translate into exponentially large increases in spontaneous Ca release frequency. This nonlinearity also works in the opposite direction, so that modest improvements in cluster integrity could produce dramatic reductions in Ca leak. These results shows that structural stability itself is a valuable therapeutic target. Strategies that preserve or restore RyR2 subunit coupling, such as enhancing the stabilizing action of accessory proteins like FKBP12.6 (38), offer a direct way to reinforce this stability. Other proteins that regulate RyR2 positioning or inter-channel contacts could likewise be harnessed to maintain cluster integrity (39). Such approaches differ fundamentally from traditional interventions aimed at modifying channel gating kinetics or expression levels, which may have limited efficacy if the underlying cluster architecture remains compromised. By identifying inter-subunit coupling as a central determinant of spontaneous Ca release, our study highlights a specific and tunable structural feature of the RyR2 complex that could be targeted to suppress pathological Ca leak.

### Model Limitations

Our model has several important limitations. First, we do not incorporate the complex regulatory mechanisms known to modulate RyR2 gating in cardiac myocytes, including inhibition by physiological Mg concentrations and modulation by regulatory proteins such as CaMKII, calmodulin, and FKBP12.6 (17,35,36,40). These factors will alter the spark waiting times. However, they are unlikely to eliminate the exponential dependence on cluster connectivity that emerges from cooperative gating between channels. Second, we model RyR2 clusters using idealized geometric arrangements of either pure oblique or pure adjoining configurations, whereas native clusters in cardiac cells exhibit heterogeneous combinations of both interaction types. Our results indicate that mixed geometries will yield intermediate waiting times, with the precise value depending on the relative proportion of each interaction type. However, because both pure configurations exhibit exponential sensitivity to coupling strength and connectivity, we expect this relationship to persist regardless of geometric details. Third, our framework does not include channel inactivation, SR Ca depletion, or competing Ca clearance mechanisms such as the sodium calcium exchanger and the sarcoplasmic reticulum Ca ATPase (SERCA). These processes determine spark termination and recovery. However, they occur on timescales much shorter than the diastolic waiting times we investigate and therefore do not affect the spark initiation dynamics that are the focus of this study. While future work incorporating more detailed RyR2 kinetics and realistic cluster geometries will refine quantitative predictions, the central finding that spontaneous Ca spark frequency is exponentially sensitive to cluster structural integrity follows directly from the cooperativity between RyR2 channels and should remain robust.

## Author contributions

AN and YS designed research; performed research; contributed analytic tools; analyzed data and wrote the paper.

## Acknowledgements

This work was supported by the National Institute of General Medical Sciences (Award Number: 1R16GM153647-01 to YS) and the National Science Foundation (Award Number: 2320846 to YS). The funders had no role in study design, data collection and analysis, decision to publish, or preparation of the manuscript. No authors received a salary from any of these funding sources. We thank these agencies for their generous support.

## References

1. Bers, D. M. 2002. Cardiac excitation–contraction coupling. Nature. 415(6868):198–205.

2. Sheard, T. M., M. E. Hurley, A. J. Smith, J. Colyer, E. White, and I. Jayasinghe. 2022. Threedimensional visualization of the cardiac ryanodine receptor clusters and the molecular-scale fraying of dyads. Philosophical Transactions of the Royal Society B. 377(1864):20210316.

3. Hurley, M. E., E. White, T. M. Sheard, D. Steele, and I. Jayasinghe. 2023. Correlative superresolution analysis of cardiac calcium sparks and their molecular origins in health and disease. Open Biology. 13(5):230045.

4. Mesa, M. H., J. van den Brink, W. E. Louch, K. J. McCabe, and P. Rangamani. 2022. Nanoscale organization of ryanodine receptor distribution and phosphorylation pattern determines the dynamics of calcium sparks. Biophysical Journal. 121(3):378a.

5. Baddeley, D., I. Jayasinghe, L. Lam, S. Rossberger, M. B. Cannell, and C. Soeller. 2009. Optical single-channel resolution imaging of the ryanodine receptor distribution in rat cardiac myocytes. Proceedings of the National Academy of Sciences. 106(52):22275–22280.

6. Kolstad, T. R., J. van den Brink, N. MacQuaide, P. K. Lunde, M. Frisk, J. M. Aronsen, E. S. Norden, A. Cataliotti, I. Sjaastad, and O. M. Sejersted. 2018. Ryanodine receptor dispersion disrupts Ca2+ release in failing cardiac myocytes. Elife. 7:e39427.

7. Soeller, C., and I. D. Jayasinghe. 2018. Quantitative Super-Resolution Microscopy of Cardiomyocytes. In Microscopy of the Heart. Springer, pp. 37–73.

8. Jayasinghe, I., M. B. Cannell, and C. Soeller. 2009. Organization of ryanodine receptors, transverse tubules, and sodium-calcium exchanger in rat myocytes. Biophysical Journal. 97(10):2664–2673.

9. Fowler, E. D., and S. Zissimopoulos. 2022. Molecular, subcellular, and arrhythmogenic mechanisms in genetic RyR2 disease. Biomolecules. 12(8):1030.

10. Waddell, H. M., V. Mereacre, F. J. Alvarado, and M. L. Munro. 2023. Clustering properties of the cardiac ryanodine receptor in health and heart failure. Journal of molecular and cellular cardiology. 185:38–49.

11. Macquaide, N., H.-T. M. Tuan, J.-i. Hotta, W. Sempels, I. Lenaerts, P. Holemans, J. Hofkens, M. S. Jafri, R. Willems, and K. R. Sipido. 2015. Ryanodine receptor cluster fragmentation and redistribution in persistent atrial fibrillation enhance calcium release. Cardiovascular research. 108(3):387–398.

12. Dixon, R. E. 2022. Nanoscale organization, regulation, and dynamic reorganization of cardiac calcium channels. Frontiers in physiology. 12:810408.

13. Benitah, J.-P., R. Perrier, J.-J. Mercadier, L. Pereira, and A. M. Gómez. 2021. RyR2 and calcium release in heart failure. Frontiers in physiology. 12:734210.

14. Chen-Izu, Y., C. W. Ward, W. Stark Jr, T. Banyasz, M. P. Sumandea, C. W. Balke, L. T. Izu, and X. H. Wehrens. 2007. Phosphorylation of RyR2 and shortening of RyR2 cluster spacing in spontaneously hypertensive rat with heart failure. American Journal of Physiology-Heart and Circulatory Physiology. 293(4):H2409–H2417.

15. Hiess, F., P. Detampel, C. Nolla-Colomer, A. Vallmitjana, A. Ganguly, M. Amrein, H. E. Ter Keurs, R. Benítez, L. Hove-Madsen, and S. W. Chen. 2018. Dynamic and irregular distribution of RyR2 clusters in the periphery of live ventricular myocytes. Biophysical journal. 114(2):343–354.

16. Ai, X., J. W. Curran, T. R. Shannon, D. M. Bers, and S. M. Pogwizd. 2005. Ca2+/calmodulin– dependent protein kinase modulates cardiac ryanodine receptor phosphorylation and sarcoplasmic reticulum Ca2+ leak in heart failure. Circulation research. 97(12):1314–1322.

17. Marx, S. O., and A. R. Marks. 2013. Dysfunctional ryanodine receptors in the heart: new insights into complex cardiovascular diseases. Journal of molecular and cellular cardiology. 58:225–231.

18. Cannell, M. B., C. Kong, M. Imtiaz, and D. R. Laver. 2013. Control of sarcoplasmic reticulum Ca2+ release by stochastic RyR gating within a 3D model of the cardiac dyad and importance of induction decay for CICR termination. Biophysical journal. 104(10):2149–2159.

19. Gao, Z.-X., T.-T. Li, H.-Y. Jiang, and J. He. 2023. Calcium oscillation on homogeneous and heterogeneous networks of ryanodine receptor. Physical Review E. 107(2):024402.

20. Iaparov, B. I., I. Zahradnik, A. S. Moskvin, and A. Zahradníková. 2021. In silico simulations reveal that RYR distribution affects the dynamics of calcium release in cardiac myocytes. Journal of General Physiology. 153(4):e202012685.

21. Li, T.-T., Z.-X. Gao, Z.-M. Ding, H.-Y. Jiang, and J. He. 2025. Formation and regulation of calcium sparks on a nonlinear spatial network of ryanodine receptors. Chaos: An Interdisciplinary Journal of Nonlinear Science. 35(2).

22. Cabra, V., T. Murayama, and M. Samsó. 2016. Ultrastructural analysis of self-associated RyR2s. Biophysical journal. 110(12):2651–2662.

23. Woll, K. A., and F. Van Petegem. 2022. Calcium-release channels: structure and function of IP3 receptors and ryanodine receptors. Physiological Reviews. 102(1):209–268.

24. Greene, D. A., T. Luchko, and Y. Shiferaw. 2023. The role of subunit cooperativity on ryanodine receptor 2 calcium signaling. Biophysical Journal. 122(1):215–229.

25. Hou, Y., I. Jayasinghe, D. J. Crossman, D. Baddeley, and C. Soeller. 2015. Nanoscale analysis of ryanodine receptor clusters in dyadic couplings of rat cardiac myocytes. Journal of molecular and cellular cardiology. 80:45–55.

26. Galice, S., Y. Xie, Y. Yang, D. Sato, and D. M. Bers. 2018. Size matters: ryanodine receptor cluster size affects arrhythmogenic sarcoplasmic reticulum calcium release. Journal of the American Heart Association. 7(13):e008724.

27. Bers, D. M., and A. Peskoff. 1991. Diffusion around a cardiac calcium channel and the role of surface bound calcium. Biophysical journal. 59(3):703–721.

28. Swietach, P., K. W. Spitzer, and R. D. Vaughan-Jones. 2010. Modeling calcium waves in cardiac myocytes: importance of calcium diffusion. Frontiers in bioscience (Landmark edition). 15:661.

29. Langer, G., and A. Peskoff. 1996. Calcium concentration and movement in the diadic cleft space of the cardiac ventricular cell. Biophysical journal. 70(3):1169–1182.

30. Asfaw, M., E. Alvarez-Lacalle, and Y. Shiferaw. 2013. The timing statistics of spontaneous calcium release in cardiac myocytes. PLoS One. 8(5):e62967.

31. Shen, X., J. van den Brink, A. Bergan-Dahl, T. R. Kolstad, E. S. Norden, Y. Hou, M. Laasmaa, Y. Aguilar-Sanchez, A. P. Quick, and E. K. Espe. 2022. Prolonged β-adrenergic stimulation disperses ryanodine receptor clusters in cardiomyocytes and has implications for heart failure. Elife. 11:e77725.

32. Mukherjee, S., N. L. Thomas, and A. J. Williams. 2012. A mechanistic description of gating of the human cardiac ryanodine receptor in a regulated minimal environment. Journal of General Physiology. 140(2):139–158.

33. Xu, L., and G. Meissner. 2004. Mechanism of calmodulin inhibition of cardiac sarcoplasmic reticulum Ca2+ release channel (ryanodine receptor). Biophysical journal. 86(2):797–804.

34. Laver, D., T. Baynes, and A. Dulhunty. 1997. Magnesium inhibition of ryanodine-receptor calcium channels: evidence for two independent mechanisms. The Journal of membrane biology. 156(3):213–229.

35. Walweel, K., and D. Laver. 2015. Mechanisms of SR calcium release in healthy and failing human hearts. Biophysical reviews. 7(1):33–41.

36. Zahradníková, A., J. Pavelková, M. Sabo, S. Baday, and I. Zahradník. 2025. Structure-based mechanism of RyR channel operation by calcium and magnesium ions. PLOS Computational Biology. 21(4):e1012950.

37. Gillespie, D. T. 2007. Stochastic simulation of chemical kinetics. Annu. Rev. Phys. Chem. 58(1):35–55.

38. Wehrens, X. H., S. E. Lehnart, F. Huang, J. A. Vest, S. R. Reiken, P. J. Mohler, J. Sun, S. Guatimosim, L.-S. Song, and N. Rosemblit. 2003. FKBP12. 6 deficiency and defective calcium release channel (ryanodine receptor) function linked to exercise-induced sudden cardiac death. Cell. 113(7):829–840.

39. Reynolds, J. O., A. P. Quick, Q. Wang, D. L. Beavers, L. E. Philippen, J. Showell, G. Barreto-Torres, D. J. Thuerauf, S. Doroudgar, and C. C. Glembotski. 2016. Junctophilin-2 gene therapy rescues heart failure by normalizing RyR2-mediated Ca2+ release. International journal of cardiology. 225:371–380.

40. Rokita, A. G., and M. E. Anderson. 2012. New therapeutic targets in cardiology: arrhythmias and Ca2+/calmodulin-dependent kinase II (CaMKII). Circulation. 126(17):2125–2139.

